# Zoogeographic patterns on very small spatial scales in rock-dwelling *Plectostoma* snails from Borneo (Gastropoda: Caenogastropoda: Diplommatinidae)

**DOI:** 10.1101/2022.02.13.480235

**Authors:** Menno Schilthuizen, Angelique van Til

## Abstract

We mapped the fine-grained distribution patterns of *Plectostoma* microsnails on two small isolated limestone outcrops in Malaysian Borneo. On both hills, two species were present (*P. simplex* and *P. concinnum* on Tandu Batu; *P. fraternum* and *P. concinnum* on Tomanggong Besar), but the patterns we found were different. On Tomanggong Besar, the two species occupy different parts of the hill and meet along a narrow hybrid zone that is characterised by a significantly higher rate of predation by *Atopos* slugs. On Tandu Batu, the two species broadly overlap and do not form hybrids. The predation rate here is the same in both species, regardless of whether they occur in monospecific localities or in mixed localities. Our results show that even small limestone outcrops of a few hundred m in diameter cannot be considered to be uniformly populated by limestone dwelling snails, and a detailed analysis of intra-hill patterns may reveal information on species differences and maintenance.

## Introduction

In elucidating the evolutionary history of species, it is often instructive to analyse geographical patterns of distribution and interaction of closely related species. Such patterns can teach us about the processes that drive and maintain species differences, their coexistence, and reproductive isolation. For example, in sympatric species that are vicariant on a small spatial scale (e.g., the central-European species of *Nicrophorus* buying beetles; Pukowksi, 1933) competition may have driven habitat differentiation. On the other hand, sympatric, ecologically similar species that are not vicariant, may still be experiencing active competition, or may have differentiated ecologically in other, non-geographic ways (e.g. *Yponomeuta* moths specialised on different host plants; Menken, 1996). An even richer image may be obtained when reproductive interactions are included, for example in hybrid zones, i.e., areas where related species meet, mate, and produce hybrids (Barton & Hewitt, 1985; Szymura & Barton, 1986; Harrison, 1993). The power of hybrid zone analysis is particularly strong if multiple zones can be studied for the same species complex (Szymura & Barton, 1991; Harrison, 1993; Arntzen et al., 2017). Such studies have given us insights into: the number of genes involved in reproductive isolation, the strength of selection on each of those genes, the presence or absence of interactions with the environment, the age of the reproductive interaction, whether of not the hybrid zones are primary or secondary in origin (and, therefore, whether the speciation process has been allopatric or not), and their eventual fate. In other words, hybrid zones are ‘natural laboratories’ where the genetic, evolutionary, and ecological forces that drive speciation and maintain species differences can be studied in the field.

Although the literature on hybrid zones has, for historical reasons, been dominated by vertebrates, land snails are particularly suitable organisms for applying the hybrid zone approach. This has to do with their exceedingly low dispersal rate, which results in evolutionary patterns in very small geographic areas (Schilthuizen, 2002). Endemism, parapatric distribution patterns, clinal geographic variation, and hybrid zones in land snails are often displayed on spatial scales of kilometres or less (Schilthuizen, 1994; Haase et al., 2013; Stankowksi et al., 2020). This means that in land snail studies, an individual researcher can, in one field day, literally walk through evolutionary patterns which, in many other organisms, would be displayed only at the scale of entire continents. For example, the analysis of five areas of contact between different members of the *Albinaria hippolyti* species complex in a 25 x 35 km region in central Crete (Schilthuizen & Lombaerts, 1995; Schilthuizen, 1995; Schilthuizen et al., 1999; Lammers et al., 2013), revealed that the clines in traits are 20-260 m wide, are usually not associated with environmental features, and involve weak selection against large numbers of genes (on the order of tens to hundreds). In another European clausiliid genus, *Alopia*, Koch *et al.* (2020) revealed the roles of coiling reversal, sexual selection, and random genetic drift in an entire non-adaptive evolutionary radiation of nine foms (species and lower taxonomic units) within an area in the Southern Carpathians of just a few kilometres across. And within a single square kilometre on Rosemary Island off Northwestern Australia, Stankowski (2013) dissected the ecological speciation process by which a globose-shelled *Rhagada* species has evolved into a dramatically different flat-shelled form across an ecological transition from a vegetated to a rocky habitat. The distances over which this transition takes place in some places are just a few tens of metres.

Although the spatial scale is very small, most of the above-mentioned land snail studies take place within a more or less extensive and continuous area of suitable habitat. The situation becomes more complex when the suitable habitat, and therefore the snail populations, are themselves fragmented and patchy. In many parts of Southeast Asia, limestone outcrops exist as small fragments of exposed karst within a “sea” of other types of substratum. In most parts of Borneo, for example, limestone outcrops are each just a few hundred metres across, but are separated from other outcrops by tens of kilometres of sandstone or mudstone. It is known (Clements et al., 2006) that limestone offers very suitable environmental conditions for calcium-dependent organisms such as land snails. Nonetheless, most land snail taxa can also occur, albeit at lower densities, on non-limestone soils.

However, there are a few snail taxa that are strictly obligate limestone-dwellers. In Borneo, the most extreme example of this is the genus *Plectostoma* Adams, 1865, which are purely restricted to the limestone outcrops on the island (Schilthuizen et al., 2003a). The concomitant geographical isolation, local adaptation, and rarity of long-distance dispersal has resulted in high numbers of, often locally endemic, species (Vermeulen, 1991, 1994).

In the Lower Kinabatangan River Valley of Sabah, Malaysian Borneo, Schilthuizen et al. (2006) and Hendriks et al. (2019) studied a system of around 20 isolated limestone hills within a region of 10 x 35 km. These outcrops are on average half a km in diameter and separated from the nearest outcrop by about 5 km of unsuitable habitat. The dominent *Plectostoma* clade in this area is the *P. concinnum* complex. Molecular phylogenetic analysis of this complex (Schilthuizen et al., 2006) has shown that it consists of the paraphyletic, highly variable species *P. concinnum* (Fulton, 1901), as well as several short-range endemic species with distinct morphological autapomorphies, viz., *P. simplex* (Fulton, 1901), *P. mirabile* (Smith, 1893), and *P. fraternum* (Smith, 1905) (Vermeulen & Liew, 2022). Most of the limestone hills in the Lower Kinabatangan Valley are inhabited by a single form from the *Plectostoma concinnum* species complex, but three hills are home to two separate forms of the species complex. Each of all these populations is distinct genetically and in shell shape and ornamentation. Genetic analysis revealed that it is an old (several million years) radiation and that the allopatric speciation process involves both genetic drift and local adaptation, primarily in response to predation by *Atopos rapax* (Rathouissidae), a molluscivorous slug (Liew & Schilthuizen, 2014; Vermeulen & Liew, 2022).

Here, we focus on the microgeographical situation on two of those hills where two species from this complex occurred sympatrically: Tandu Batu Hill (*P. simplex* and *P. concinnum)* and Tomanggong Besar (*P. fraternum* and *P. concinnum).* Genetic studies have already shown that in neither of these cases, the two sympatric species are each other’s closest relatives, which means that each of these hills must have been colonised twice independently. In this paper, we map the geographic distribution of the sympatric species on both hills in detail, and we also assess the presence of absence of hybridisation. Finally, we investigate to what extent the species differ in suffering from slug predation.

## Materials and Methods

The two hills (Tandu Batu, 118°20’34.3”E 5°35’47.5”N, and Tomanggong Besar, 118°18’17.3”E 5°30’56.3”N) were visited from October to December, 2003. On both hills, 45 localities of 5 x 5 m were chosen that lay within suitable *Plectostoma* habitat (exposed vertical limestone rock surfaces) but otherwise were randomly distributed over the hill. At each locality, coordinates were recorded with a hand-held GPS device (Garmin), and 50 adult *Plectostoma* individuals were removed from the rock and placed into a 2.0 mL vial with 100% ethanol, and labelled. In the laboratory, each individual was identified to species. Individuals with clearly intermediate morphology were assumed to be hybrids and separately recorded.

In addition, from each locality, a 5 L soil sample was taken from underneath all vertical limestone surfaces. The soil was sieved with 5 mm mesh width, and the sieved fraction stored in a labelled plastic bag. In the laboratory, empty adult *Plectostoma* shells were extracted from these soil samples by placing the soil in water, removing all floating debris, drying this, and then passing it under a dissecion microscope to pick out the shells (Tweedie, 1961). The shell of each individual was identified to species, and the typical holes produced during predation by *Atopos rapax* slugs (Schilthuizen et al., 2003b, 2006; Liew & Schilthuizen, 2014) were recorded. Again, individuals with clearly intermediate morphology were marked as hybrids and separately recorded.

Besides taking samples, we also recorded any absence of *Plectostoma* species as well as microhabitat features such as limestone boulders and cliffs (or the absence of bare rock) while traversing the two sites.

## Results

### Distribution patterns

On Tomanggong Besar (Fig. 1), we found that most (ca. 75%; 27 localities) of the hill is occupied by a small-shelled population of *P. concinnum.* Only the northwestern flank of the hill, as well as the small ‘satellite’ hillock, are occupied by the larger-shelled *P. fraternum* (14 localities). Unexpectedly, in one locality (locality 1, on the far western side of the hill, close to the river’s edge) among *P. concinnum*, we also found several individuals of *P. brevituba* (Vermeulen, 1994), which is not closely related to the *P. concinnum* complex. The distribution areas of *P. concinnum* and *P. fraternum* adjoin along a zone that mostly coincides with a steep limestone cliff of 5-10 m in height. At four localities that lie along this zone, no pure individuals of either species were found; instead these localities are populated by what appear to be individuals that are genetic mixtures (primary and secondary hybrids as well as back-crosses) of both species. On the summit of the hill, very little suitable habitat is available, and *Plectostoma* is absent or very scarce across most of this area.

**1.**
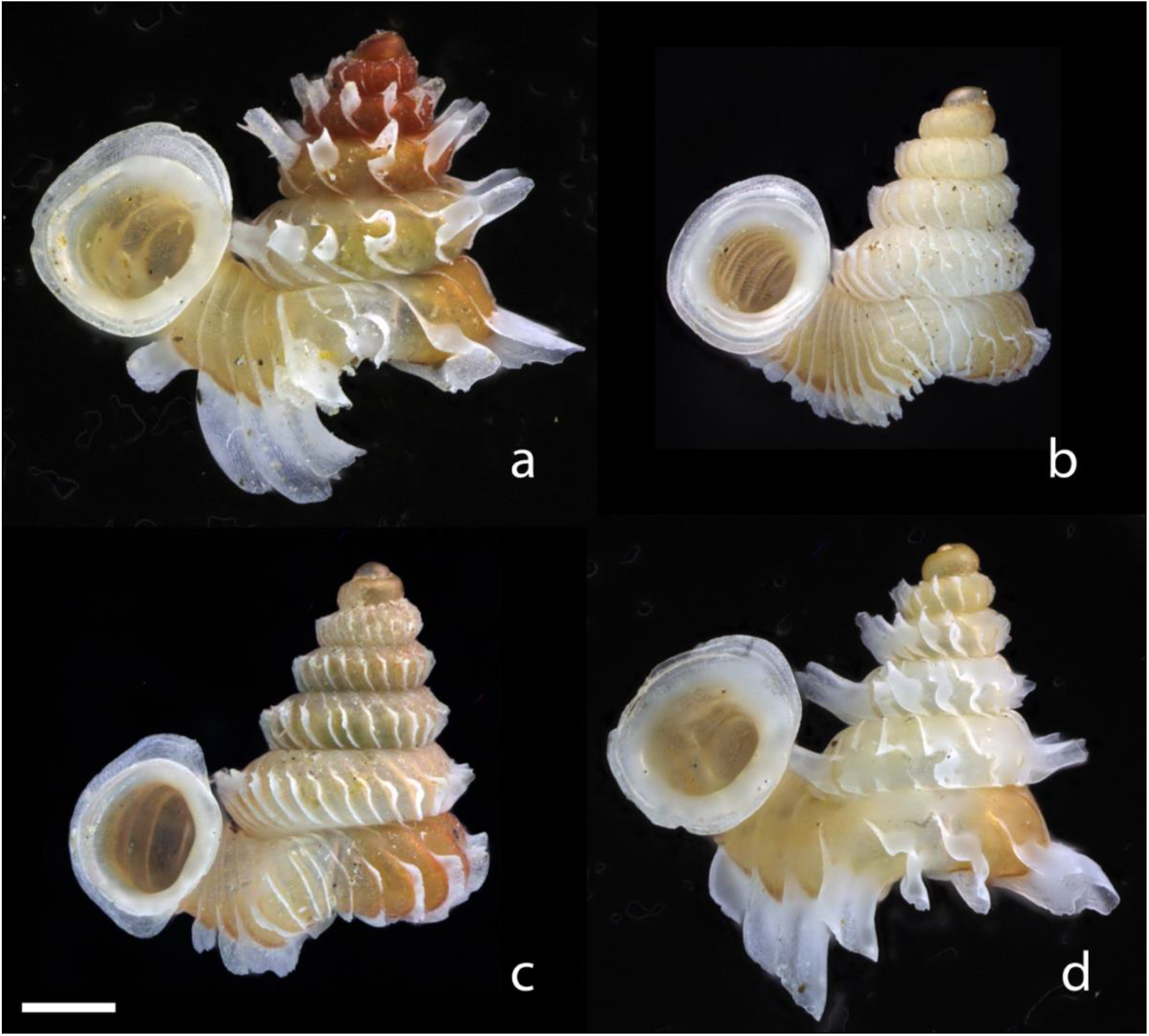
Photos of shells of (a) *P. fraternum* (Tomanggong Besar), (b) *P. concinnum* (Tomanggong Besar), (c) *P. simplex* (Tandu Batu), and (d) *P. concinnum* (Tandu Batu). Scale bar = 0.5 mm

On Tandu Batu, the situation is different (Fig. 2). Here, *P. concinnum* is mostly found on the northwestern side of the hill (14 localities), *P. simplex* (18 localities) mostly on the southeastern side (as well as on the small satellite hillock to the south), but in between and scattered throughout the hill are many (13) localities where both species co-occur, without any sign of hybridization. In these mixed populations, *P. simplex* was the most abundant overall.

**2.**
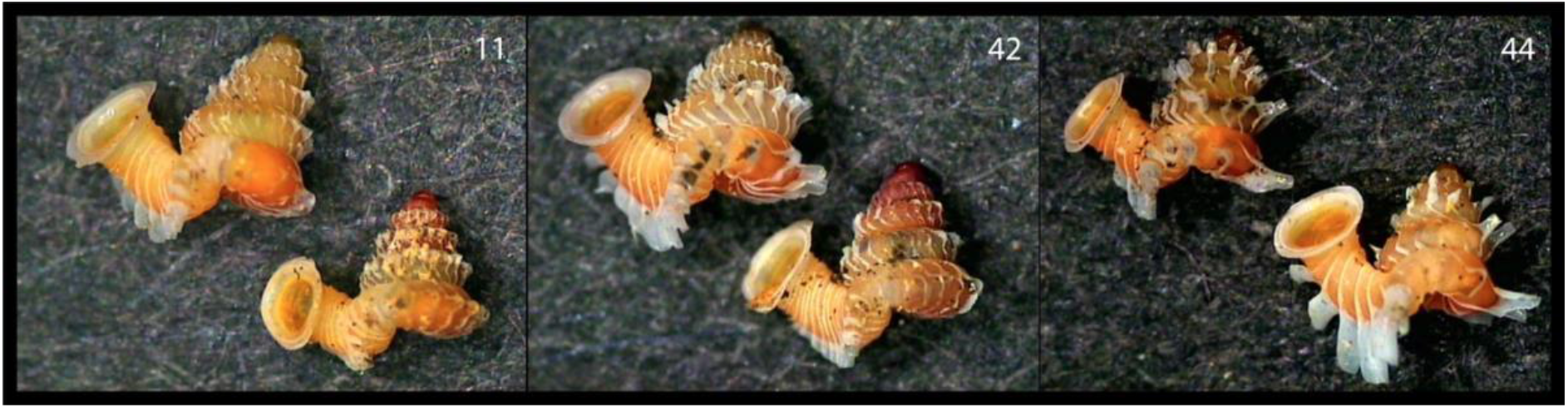
Photos of morphologically intermediate shells from the *P. concinnum* x *P. fraternum* hybrid populations 11, 42, and 44 on Tomanggong Besar (images taken from Van Til, 2004).

**3.**
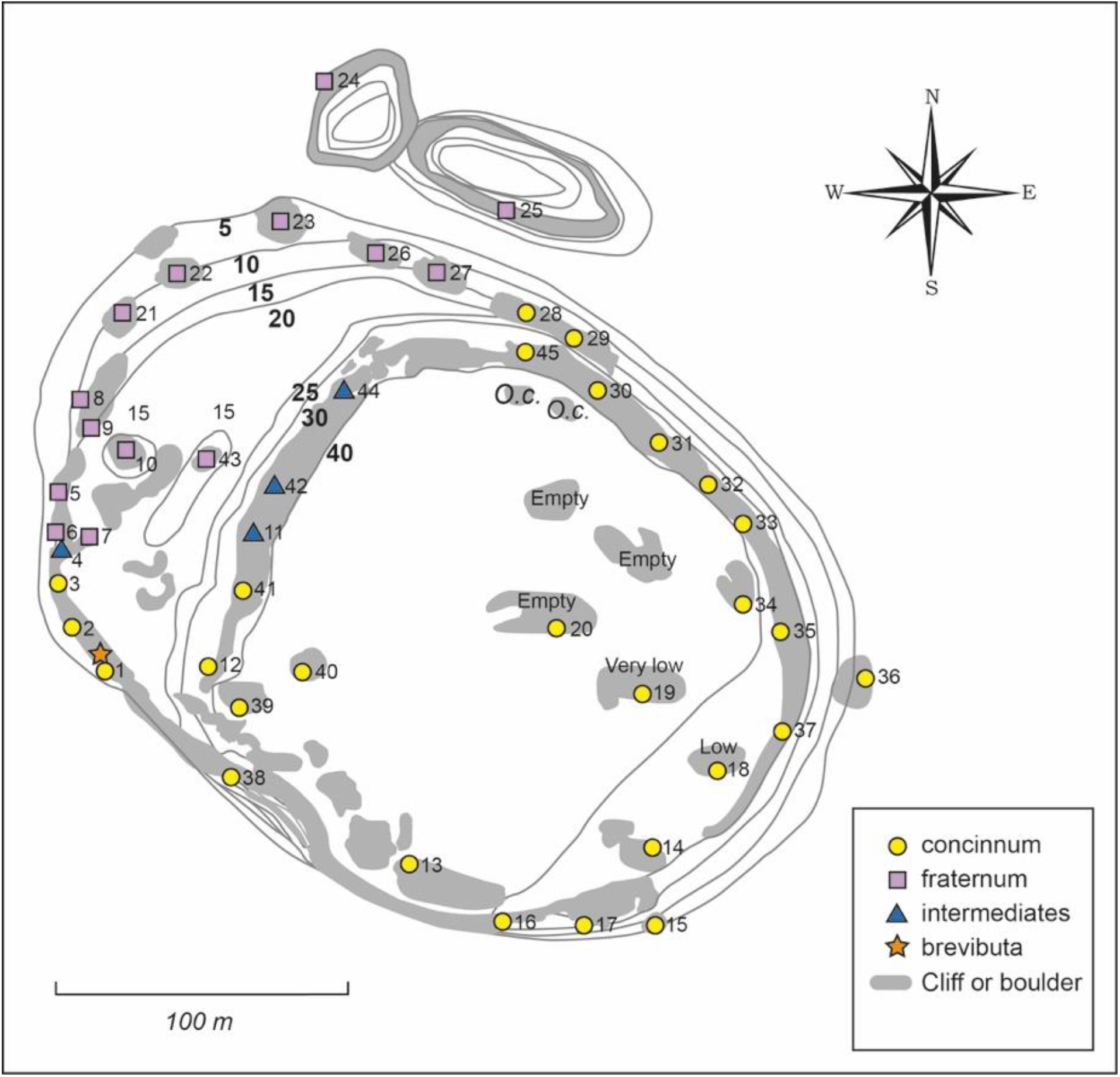
Distribution map of *P. concinnum, P. fraternum, P. brevituba*, and *concinnum* x *fraternum* hybrid populations on Tomanggong Besar.

**4.**
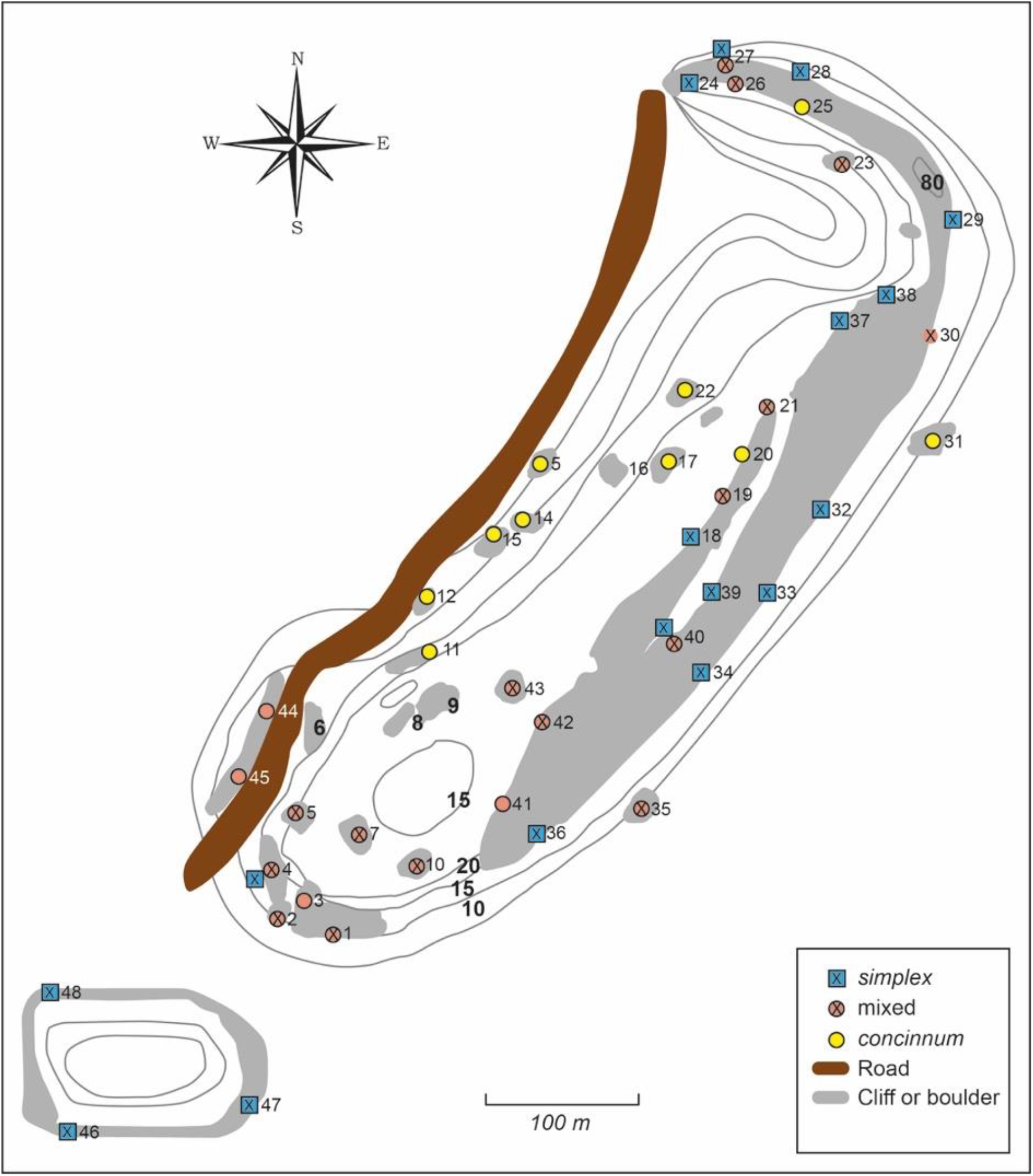
Distribution map of *P. concinnum* and *P. simplex* on Tandu Batu

The above results are based only on the living population. Looking at the samples of empty shells from the forest floor (which may include specimens from decades ago; Schilthuizen, 2011), we found the following regarding distributions. On Tomanggong Besar, the dead shells showed the same distributions as the living population. On Tandu Batu, there were several localities where the living population consisted entirely of *P. concinnum*, whereas the shells in the soil showed a mixture of both species.

### Predation

The patterns of predation are based on the empty shells taken from the forest floor and may therefore be an average across multiple years. On Tomanggong Besar, we found a mean *Atopos rapax* predation rate of 0.17. The rate in *P. fraternum* was significantly higher than in *P. concinnum* (0.19 vs. 0.14, respectively; *P* < 0.001; Chi-square test). Interestingly, the hybrids showed a significantly higher predation rate than either of the pure populations, namely 0.26 (*P* = 0.002; Chi-square test). On Tandu Batu, the mean *Atopos* predation rate was 0.16. The rate in *P. simplex* was somewhat lower than in *P. concinnum* but not statistically significantly so (0.12 vs. 0.16, respectively; *P* = 0.16; Chi-square test). In the mixed localities, predation rate was 0.17; this was also not significantly different from the predation rates in either of the monospecific populations. Although predation rates on both hills vary greatly between localities and, in the case of Tomanggong Besar, between *Plectostoma* species, there were no indications of further geographic patterns in predation rate across each of the two hills.

## Discussion

Even for snails, the geographical patterns that we have revealed in this study are on the small end of the scale. This is especially the case for Tomanggong Besar, where the endemic *P. fraternum* occupies its global territory (at an estimated population size of half a million; Schilthuizen et al., 2003b), an area of approximately 200 x 75 m, and even manages to maintain a 100 m long, 10 m wide hybrid zone with the more widely distributed *P. concinnum.* The fact that the hybrid zone appears to be ‘trapped’ at a geographical feature (a cliff) and is characterised by high rates of predation, suggest that it is a so-called ‘tension zone’ (Barton & Hewitt, 1985): a hybrid zone that is maintained by a balance between reduced hybrid fitness (which removes hybrid gene combinations from the zone) and dispersal (which generates new hybrid gene combinations in the zone).

We previously (Schilthuizen et al., 2006) found that shell ornamentation in *Plectostoma* may be engaged in an evolutionary arms race with the attack behaviour of *Atopos* slugs. This could explain why the hybrid population suffers greater predation rate: its shell shape is composed of elements of both parental species, which may render the hybrids imperfectly defended. It cannot be predicted whether the zone will be stable in the long run, but given that the population size and area of *P. concinnum* are greater, it is conceivable that the zone will eventually move at the expense of *P. fraternum* and the latter species will go extinct.

Interestingly, the situation on Tandu Batu is different. Here, *P. simplex* and *P. concinnum* appear to be partly parapatric, partly sympatric, while showing very little, if any, hybridization. The difference in hybridization rate with Tomanggong Besar cannot only be due to different degrees of relatedness, because both species pairs have similar phylogenetic distances (Schilthuizen et al., 2006). Another difference with the situation on Tomanggong Besar is that the predation rates across the different taxa are not distinguishable on Tandu Batu, suggesting that both species are equally well defended against *Atopos* attack.

In summary, our results show that, even on small limestone hills, microgeographic patterns in distribution, hybridization, and predation can be discerned that reveal aspects of the evolution and maintenance of species differences. This may be of relevance for the many zoologists working on the malacofauna of these small, isolated limestone outcrops. Often, especially for small hills of just a few hundred m in diameter, the assumption is that a single large sample taken on a hill is sufficient to represent the entire population of a species. Our results show that this simplified approach would leave interesting information unaccessed.

## Acknowledgements

We thank Liew Thor-Seng, Noel Tawatao & Tachaini Narainan for assistance and company in the lab in UMS. Isabel Acrenaz-Lackman, Marc Acrenaz, Berjaya Elahan, Mislyn Elahan, and Ramlan Sakong arranged for accommodation at and assistance from the Kinabatangan Orangutan Conservation Project (KOCP) in Sukau. In the field, help and support was provided by Heather C. Leasor and by many KOCP field assistants, i.e., Marli Suali, Zolirwan Takasi, Hasbollah Sinyor, Abd. Rajak Saharon, Jamil Sinyor, Eddy Ahmad, Megga Batadon, Ahmad Kapar, Hadrin Lias, Azmey Sakong, Suali Adrari, Berjaya Elahan, Adam Malan, Azlie Etin, Salleh Saharon, Rosdi Bahtar, Sulaiman Ismail, Rosman Sakong, Ramlan Sakong, Suhailie Bahar, Herman Suali, Azman Sakong, Johri Bakri, Sophia Hakim, Eddy Taib, Ahmadia Lias, and Fadlee Indal. The people of Kampung Sukau were very welcoming during our field work. In the Netherlands, encouragement and assistance was provided by Sandrine Ulenberg, Janine Mariën, Nico de With, Aad Lindeman, and Jodi Apeldoorn. This work was financially supported by an internal grant from Universiti Malaysia Sabah to M.S.. The Sabah Wildlife Department, the Sabah Forestry Department, and the Village Heads of Sukau gave permission to conduct the fieldwork.

---

**Table 1.** Predation rates on both hills. For Tandu Batu, under ‘allopatric’ all samples were combined where each species was found in isolation, under ‘sympatric’ all samples where both species were found. Predation was determined by the presence of a typical hole made by *Atopos rapax* in the shell of the prey. For Tomanggong Besar, under *P. concinnum* and *P. fraternum*, all shells were combined that could unambiguously be assigned to these species, respectively; under ‘hybrids’ all morphologically intermediate shells were combined. Different letters denote predation rates that are significantly different (within their own locality); see Results for details.

| **Tandu Batu** | **total** | **predation** | **predation rate** |
| --- | --- | --- | --- |
| *P. concinnum* allopatric | 150 | 18 | 0.12<sup>a</sup> |
| *P. simplex* allopatric | 1050 | 173 | 0.16<sup>a</sup> |
| sympatric | 1077 | 181 | 0.17<sup>a</sup> |
| **Tomanggong Besar** |  |  |  |
| *P. concinnum* | 2969 | 423 | 0.14<sup>a</sup> |
| *P. fraternum* | 1495 | 287 | 0.19<sup>b</sup> |
| hybrids | 429 | 111 | 0.26<sup>c</sup> |

